# Closed-loop modulation of remote hippocampal representations with neurofeedback

**DOI:** 10.1101/2024.05.08.593085

**Authors:** Michael E. Coulter, Anna K. Gillespie, Joshua Chu, Eric L. Denovellis, Trevor T.K. Nguyen, Daniel F. Liu, Katherine Wadhwani, Baibhav Sharma, Kevin Wang, Xinyi Deng, Uri T. Eden, Caleb Kemere, Loren M. Frank

## Abstract

Humans can remember specific remote events without acting on them and influence which memories are retrieved based on internal goals. However, animal models typically present sensory cues to trigger memory retrieval and then assess retrieval based on action. Thus, it is difficult to determine whether measured neural activity patterns relate to the cue(s), the memory, or the behavior. We therefore asked whether retrieval-related neural activity could be generated in animals without cues or a behavioral report. We focused on hippocampal “place cells” which primarily represent the animal’s current location (local representations) but can also represent locations away from the animal (remote representations). We developed a neurofeedback system to reward expression of remote representations and found that rats could learn to generate specific spatial representations that often jumped directly to the experimenter-defined target location. Thus, animals can deliberately engage remote representations, enabling direct study of retrieval-related activity in the brain.

## INTRODUCTION

Remembering is distinct from acting. Humans can remember places we have been and experiences we have had without overt behavioral signs that these memories have been retrieved. Remembering can also happen in the absence of specific external retrieval cues. Humans can retrieve specific memories based on internal goals^1^, and in some cases memories can even seem to be retrieved spontaneously^2^. Finally, remembering an experience distant in place or time does not seem to require mentally traversing all intermediate places or times; instead the brain is able to mentally teleport or “jump” directly to the memory^3^. Thus, memory retrieval is a process that can be expressed in the brain separately from: (1) the decision to act based on the content of the memory, (2) specific external cues that trigger the memory, and (3) intervening experiences that separate the current state from the past event.

Memory retrieval itself is not well isolated from these co-occurring processes in current animal models and associated experimental paradigms, which limits our ability to understand the activity patterns that support retrieval. First, current approaches typically assess whether a memory has been retrieved based on a behavioral report. Widely used paradigms including contextual-fear conditioning^4–6^, the Morris Water Maze^7^, and many others, place animals in a familiar context and quantify memory based on whether the animals exhibit specific behaviors (e.g., freezing or directed navigation to a remembered goal location). Activity patterns measured in these paradigms are the result of a complex combination of sensory information processing, memory retrieval, a decision process, and action, and so it is difficult to isolate and study the retrieval process separate from other ongoing computations.

Second, current experimental approaches typically engage retrieval by presenting external cues such as those found in a particular spatial context^7–10^. These cues can strongly influence patterns of brain activity, including spiking in brain regions like the hippocampus^11^ that contribute to memory formation and retrieval^9^. Thus, representations of the current sensory inputs can be difficult to disentangle from memory-related activity.

Third, current paradigms do not require that animals retrieve internal representations associated with experiences distant in time and place. Even in cases where animals must navigate to a distant location^7^, or where behavior depends on a past trial^12^, we cannot be sure that any particular remote memory is being retrieved. Instead, animals may use alternative strategies where they construct movement vectors toward a goal or makes choices based on familiarity.

How, then, can we directly study memory retrieval in animals? Brain-machine interface (BMI) paradigms provide a powerful approach that has the potential to enable these studies. Numerous experiments have shown that subjects can learn to control a specific neural activity pattern (from single neuron spiking to population-level activity) via continuous visual or auditory feedback (e.g. moving a cursor on a screen to a goal location)^13–17^. As a result of the sensory feedback, the subject learns to incrementally change the target neural activity pattern to reach a goal state. In the context of the hippocampus, a BMI device using this approach was recently developed wherein rats learned to mentally navigate through a virtual reality environment^18^. The device updated the visual display corresponding to a location in the environment based on population activity in hippocampus and thus, provided continuous visual feedback to the animal. In this environment, rats learned to generate representations of continuous spatial trajectories from the current position to a goal location.

These previous results demonstrate that with continuous sensory input, the brain can learn to generate the next correct pattern in a sequence to achieve a goal. However, two additional features are required to enable direct study of retrieval of a spatial memory. First, the retrieval-related pattern would need to be generated in the absence of the specific (confounding) sensory input associated with the location. Second, the retrieval related pattern would need to be generated without requiring the generation of the full intervening sequence of places between the current position and the remembered location.

We therefore developed a novel, real-time neurofeedback paradigm that did not present memory-specific cues and did not require traversal of a complete mental trajectory to arrive at a retrieved location. In our paradigm, rats were rewarded for generating remote hippocampal spatial representations. We focused on hippocampal activity patterns as a substrate for retrieval, both because the hippocampus is critical for spatial and event memory retrieval for recent experiences^9^ and because hippocampal activity patterns permit the experimental identification of activity consistent with retrieval. Specifically, as an animal explores an environment, each hippocampal place cell is predominately active when the animal occupies one or more locations in an environment (the cell’s “place fields”). Sets of place cells can also “reactivate” when the animal is far outside the cells’ place fields^11,19–25^, reinstating a remote representation. These events can also include representational jumps to remote locations that would not be possible during real movement^26^, and these events are associated with activity across the brain consistent with the retrieval of a memory^24,27,28^. Further, artificial activation of a remote representation^29^ or association of the representation with reward can be sufficient to drive associated behaviors^29,30^.

Using this paradigm, we found that rats could learn to generate specific remote hippocampal spatial representations in the absence of sensory cues indicating which representations to generate. Strikingly, in most cases, these representations jumped to the target region without representing the intermediate locations between the animal and the target. Further, these representations were largely expressed in a brain state not previously associated with remote spatial activity patterns. This work establishes a model for studying spatial memory retrieval in the absence of sensory cues and specific behavioral outputs.

## RESULTS

### Closed-loop hippocampal neurofeedback

We developed a behavioral task and a real-time neurofeedback system that rewarded animals for generating specific, experimenter-chosen remote representations. Six rats were surgically implanted with tetrodes targeting hippocampus and following recovery, participated in a feedback-based task in a modified Y-maze equipped with lighted reward ports (Fig. 1a, S1a, S2a). In each of three daily sessions, rats were first visually cued to explore the left and right arms of the environment (Fig. 1a, Task phase 1). Following this exploration period, the lighted ports in the outer arms were extinguished, and the feedback portion of the session began at a central reward port away from the arms (Fig. 1a, Task phase 2).

**Figure 1.**
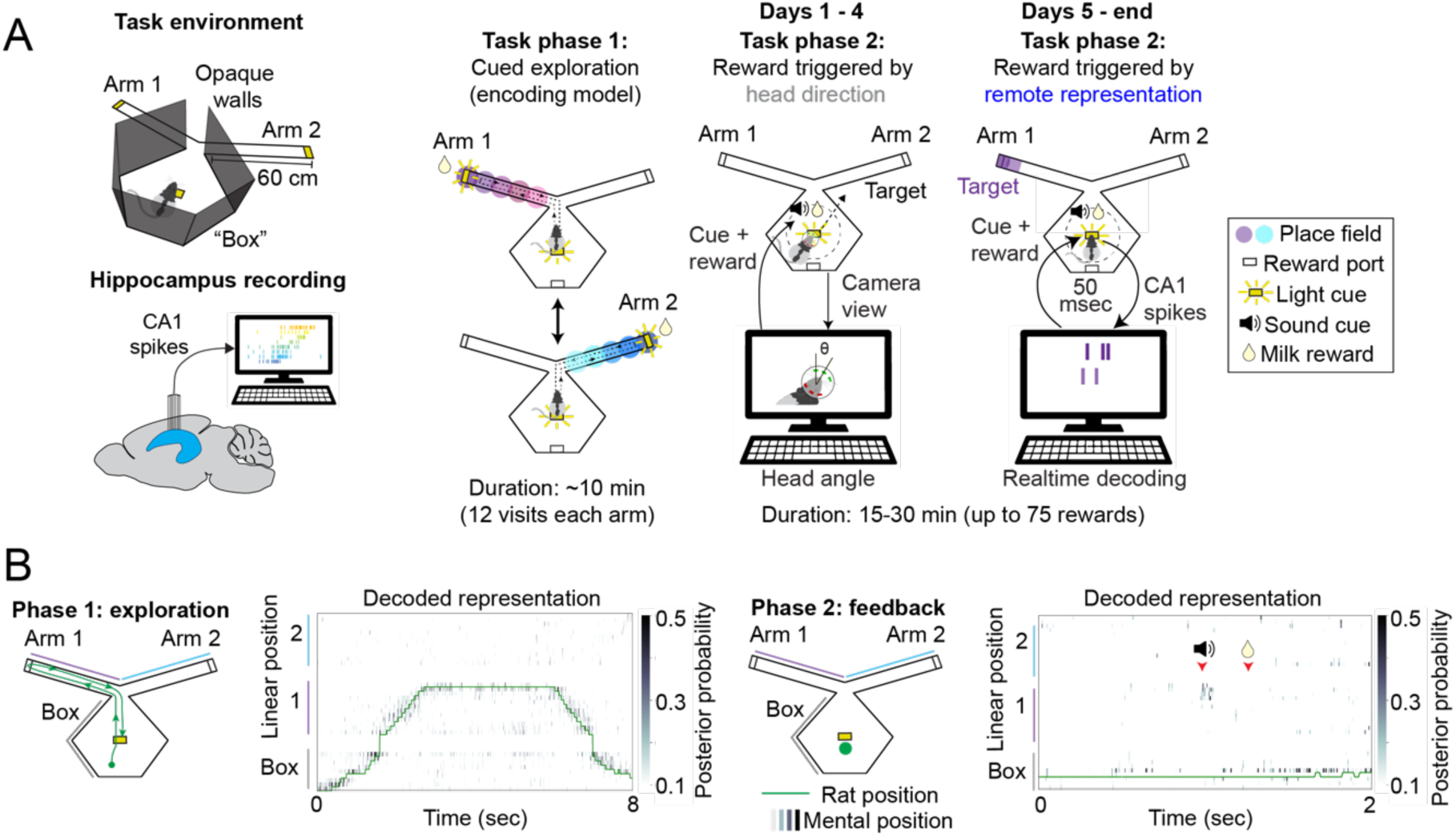
Closed-loop hippocampal feedback system. (**A**) The task environment consisted of a central “box” area with a central reward port and two arms, each with a reward port at the end. The end of one of the two arms was used as the target location for neurofeedback in each session. Note that the walls surrounding the central box are opaque. Each task session contained two task phases, exploration and feedback. During the feedback phase, either specific head directions or remote target representation were detected and triggered a tone. A nosepoke at the center well within 3 seconds of the tone then triggered delivery of reward. (**B**) Clusterless decoding of hippocampal activity accurately tracked the rat’s actual position during movement. During feedback, the decoder detected remote representations that triggered tone and reward.

The feedback portion of the task was divided into two stages. In both stages animals were required to remain near the center well of the track to receive reward. During the first stage (days 1-4 of training), rats received behavioral feedback based on head direction. If the rat turned its head in the specified direction (either left or right) and was near the central reward port, a reward-cue tone played. If the rat then nose-poked in the port within 3 sec, reward was delivered (see Methods).

Following the four days of head direction feedback, we switched to the second, neurofeedback stage. In this stage animals were rewarded only if they generated a hippocampal representation of a remote target region: the distal end of either the left or right arm of the track (see Methods and Fig. 1a). This remote representation was identified by a real-time system that continuously decoded (every 6 milliseconds) the hippocampal representation of space using an encoding model that related hippocampal spiking activity to the rat’s position during the exploration phase of each session (Fig. S1a-e)^26,31,32^. If the decoder detected a temporally extended representation of the target region the tone was presented. The animal was then rewarded if it nose-poked in the central well within 3 seconds (Fig. 1a). The target region was held fixed for three days (nine sessions) and then switched to the other arm. This switching continued for 6-18 days per animal depending on the quality of the recordings. In each session, the feedback period last for either 30 minutes or until animals received 75 rewards. Importantly, during the feedback period, the animal could not see the target location at the arm end from central port, and no target-location-specific sensory cues were presented.

Behavioral results showed that all six rats were able to rapidly and consistently maximize the number of rewards per session with behavioral feedback based on head direction (Fig. S3a). Further, all six rats also reached the maximal number of rewards per session during remote representation neurofeedback, although there was substantial variability across sessions and rats; some animals achieved high performance in many sessions while others achieved high performance in a smaller fraction of sessions (Fig. 2a).

**Figure 2.**
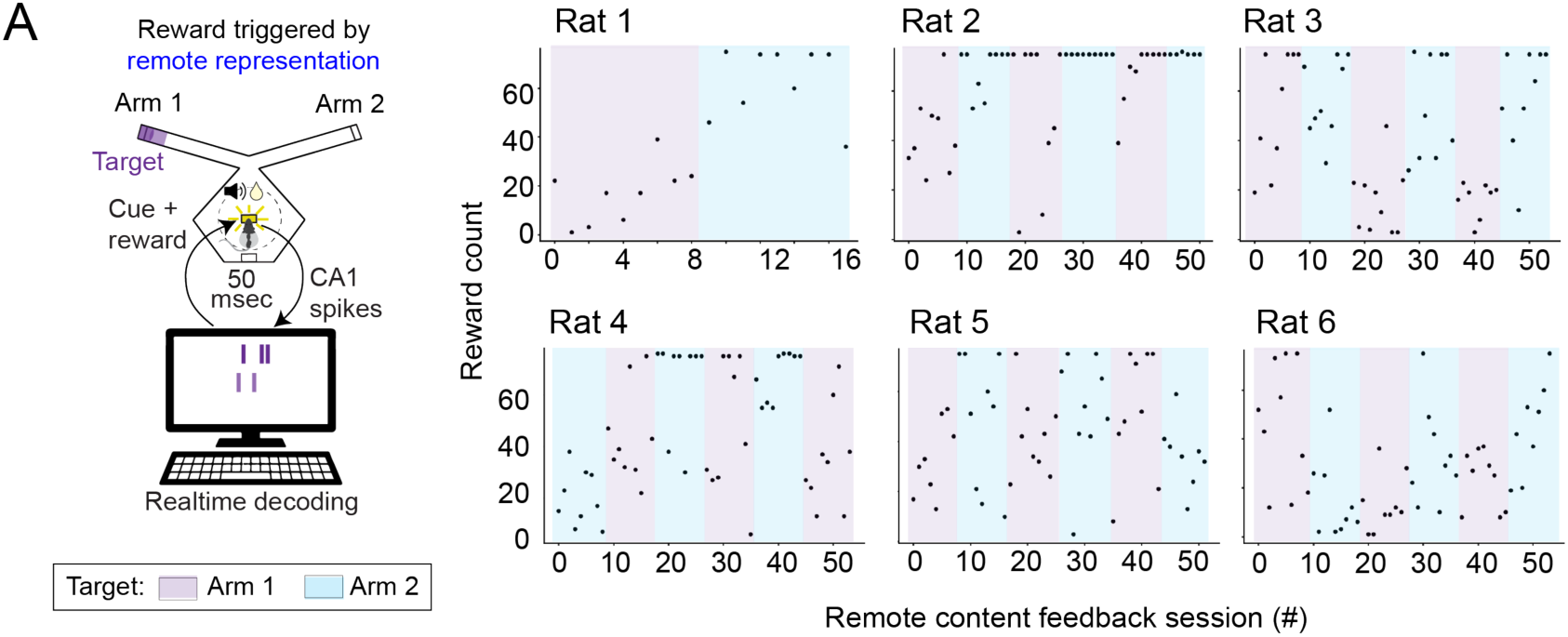
Rewards received during neurofeedback task. (**A**) Each rat maximized rewards during some neurofeedback sessions. Maximum of 75 rewards per session.

To analyze the data from the neurofeedback sessions, we first validated the accuracy of our real-time decoder in post-experimental analyses where we used the encoding model to decode the actual location of the animal during the cued exploration period (Task phase 1; Fig. 1b, left; Fig. S2b,c; Methods); a small number of lower quality decoding sessions were excluded from further analyses. We then examined the representations that triggered reward in the neurofeedback sessions to validate the real-time system. As expected, the spatial content was specific to the chosen remote target, while during head direction feedback sessions, the rat’s current location was typically represented (Fig. 1b, right, S3b,c,d).

We found that animals solved this task by generating representations that often jumped to the target region, consistent with mental teleportation, a key capacity of memory retrieval. Inspection of individual representations (up to 90 msec before detection) showed that most were confined to the target region, some included the target region and the start of the target arm, and rarely, they included partial and full remote sequences towards the target (Fig. 3a). Across all rats, 55-70% of detected representations included only the target location or jumped from the current location to the target, while only 1-4% of representations were sequential trajectories down the length of the arm (Fig. 3b). Most representations (56-73%) jumped at least 50 cm from the rat’s current location (Fig. S4a).

**Figure 3.**
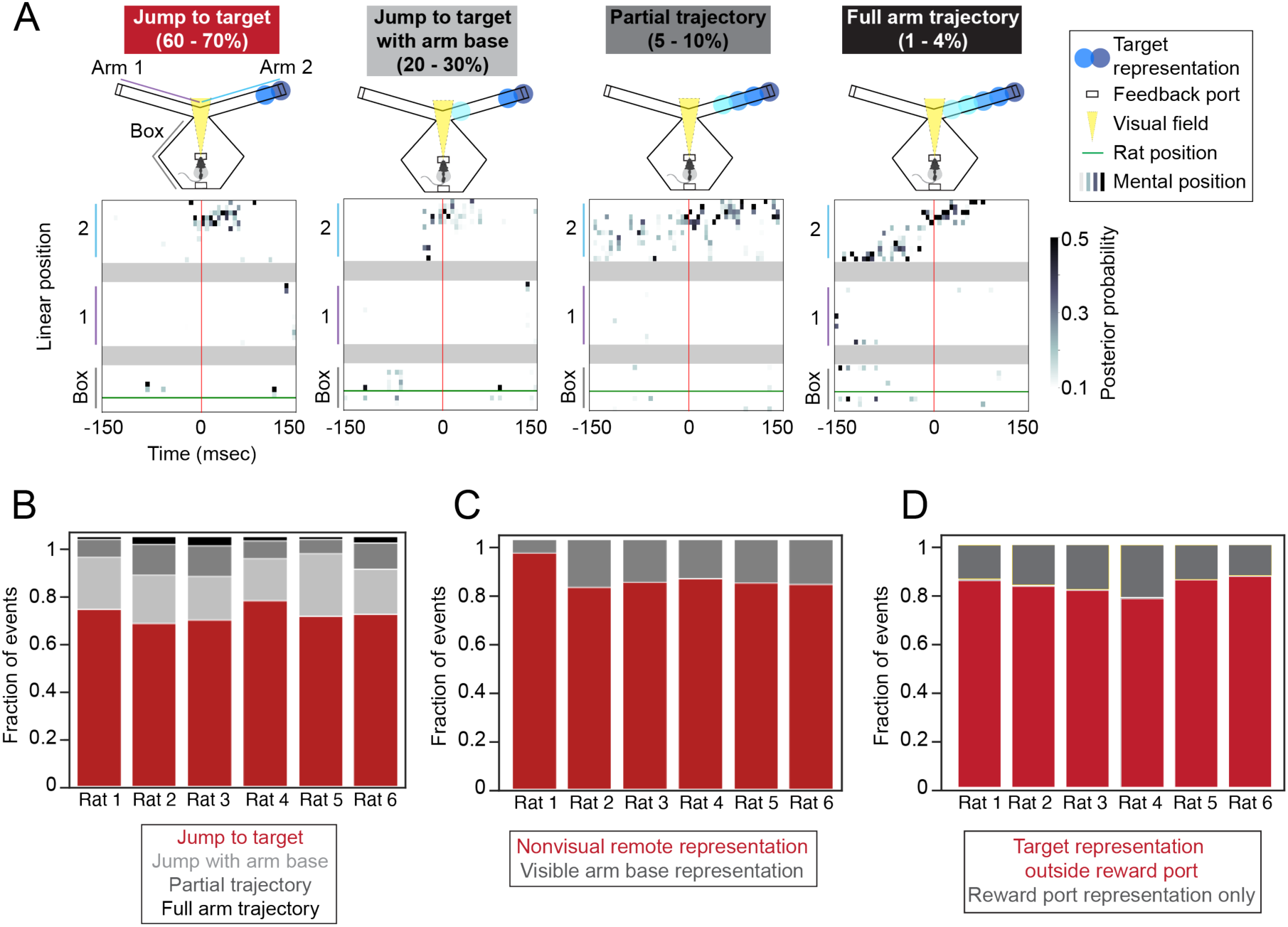
Remote representations jump to target location. (**A**) Individual detected remote representations. Representation classification and frequency are noted above each example. Colored circles on track indicate decoded location of spatial representation. (**B**) Summary of event classification for all rats. This includes 90 msec before detection. (**C**) Fraction of detected events with representation of the arm base within the visual field. 8-20% of events per rat. (**D**) Fraction of detected events with representation of the exact reward port location. 15-25% of events per rat.

Our results also suggest that neither specific sensory cues nor reward-specific representations contributed substantially to the observed remote activity patterns. First, when the animals were close to the center well as was required for reward to be delivered, they were able to see only the very beginnings of the outer arms. If this sensory input triggered retrieval, we would expect that the resulting representation would include the beginnings of the arm associated with the target location. This was not the case: these representations were present in between 6 and 20% of events across rats (Fig. 3c), suggesting any available cues were only contributing to a small fraction of events. Second, although the target location included the reward port at the very end of the arm, detected remote representations only occasionally included a representation of the reward port location (13-22% across rats; Fig. 3d), showing that generation of remote representations was not primarily driven by the target being a reward location.

### Increased representation of the target location

Our system established relatively strict criteria for a remote spatial representation (see Methods), and a detailed inspection of the decoded representations present during the neurofeedback periods revealed numerous instances of apparent target representations that did not meet these criteria. Therefore, to determine whether target representation was consistently enriched through neurofeedback (which would provide further evidence that animals can control representation content), we calculated the fraction of time that at least 40% of the representation (probability mass) was within the target region while the rat was at the center port (see Methods; Fig. 4a,b). During these periods, the majority of the representation was almost always within the target arm (Fig. S4b).

**Figure 4.**
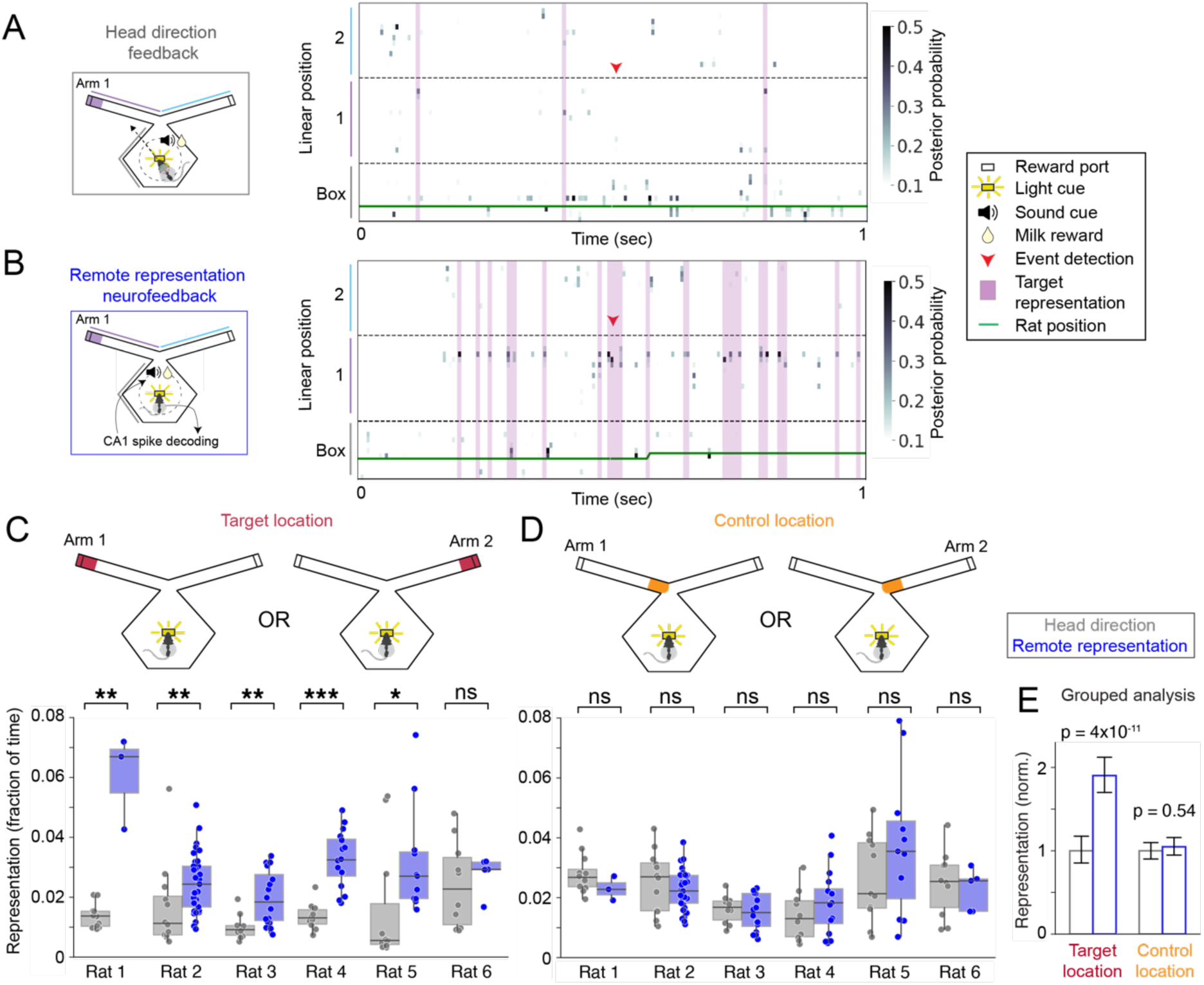
Increased hippocampal remote representations during neurofeedback. **(A)** Head direction feedback (schematic at left) and example of decoded mental position around the time of a detected correct head direction event (right). **(B)** Remote representation neurofeedback session (schematic at left) and example of decoded mental position around the time of a remote representation detection event (right). **(C)** Target location representation prevalence during high-reward sessions for head direction feedback vs remote representation feedback sessions for each rat. **(D)** Control location (non-rewarded) representation prevalence during high-reward sessions for head direction feedback vs. remote representation feedback sessions for each rat. **(E)** Grouped (LME) analysis across all 6 rats. *: p<0.05, Mann-Whitney test, **: p<0.01, ***: p<0.001.

We then compared remote representation during head direction feedback sessions to remote representation during neurofeedback sessions. Because reward itself can drive patterns of hippocampal activity that express remote representations^33,34^, we restricted our analyses to sessions with similar reward amounts (>90% of possible rewards). We found that remote representation neurofeedback drove a substantial increase in the generation of representations of the target region as compared to the head direction feedback. Representations of the target region were approximately two-fold more prevalent during neurofeedback sessions compared to head direction feedback sessions (Fig. 4c,e, individually significant in 5 of 6 rats (p < 0.05, Mann-Whitney); grouped analysis (linear mixed effects model, LME): all data: p = 3.9e-11, tone-triggering representation removed: p = 3.6e-11; Fig. S5a). Importantly, this did not reflect a non-specific increase in remote representations: representations of a physically closer location (the base of the target arm) were not more prevalent in neurofeedback sessions compared to head direction feedback sessions (Fig. 4d,e, non-significant in each rat, grouped analysis: p = 0.54, LME). Likewise, representation of the end of the opposite (non-target) arm was not enriched (Fig. S5c, grouped analysis: p = 0.067, LME). We also examined the relationship between task performance (number of rewards received) and the amount of target representation and found that these measures were significantly positively correlated in five of the six animals (p < 0.01 for rats 1, 2, 3, 5 and 6) and weakly negatively correlated in one animal (p = 0.022, rat 4).

The prevalence of these target representations increased with time, consistent with a learning process. We examined target representation across all included remote representation neurofeedback sessions (6 or 18 days) and found that the even though the target representation switched every three days, the overall prevalence of the target representation increased over time (Fig. 5, individually significant in 4 of 6 animals; grouped analysis: linear regression of z-scored values, all data: p = 0.0001, tone-triggering representation removed: p = 0.0002; Fig. S5b). By contrast, there was no consistent change in the representation of the base of the target arm (Fig. 5, significant increase in only one animal; grouped analysis: linear regression of z-scored values, p =0.7). These findings complement the analyses of content for the reward triggering events (Fig. 3b) to confirm that rats most often generated representations with discontinuous jumps from the animal’s current location to the distant target location rather than representations that moved sequentially to the target location. Finally, we also noted that representation of the opposite arm end also increased over time (Fig. S5d, linear regression of z-scored values, p = 0.004), perhaps because the target switched back and forth between the two arms every three days. Overall, the longitudinal increases in representation were specific to rewarded locations.

**Figure 5.**
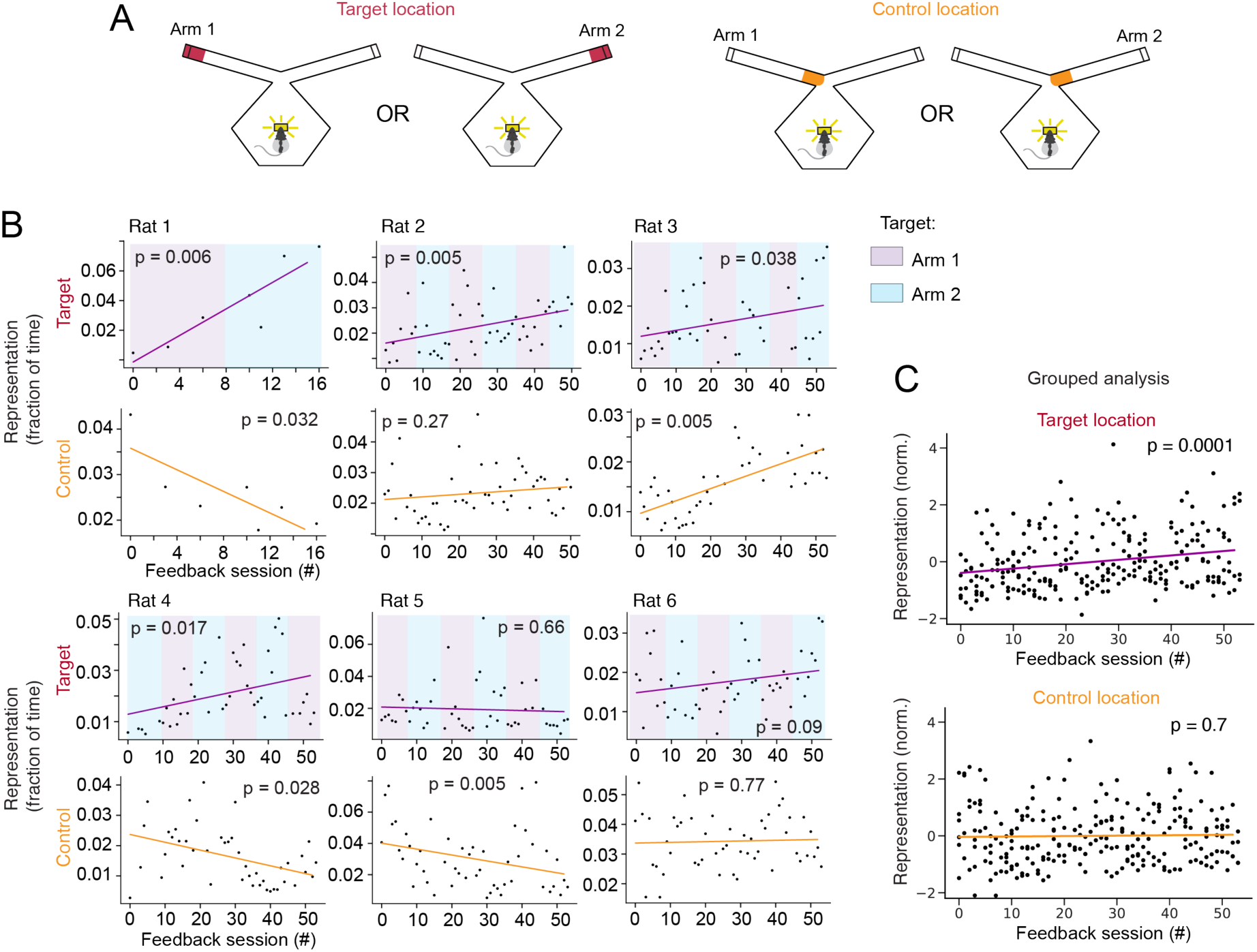
Remote hippocampal representations increase over time with neurofeedback. **(A)** Target location is the arm end and control location is the arm base. **(B)** Target location representation prevalence across all remote representation feedback sessions (red) and for control location representations (orange). Colors in top plots represent designated target arm. Line shows linear fit; p-value corresponds to the slope of the linear fit. **(C)** Grouped analysis (linear regression) for normalized data from all 6 rats for target (top) and control locations (bottom). *: p<0.05, Mann-Whitney test, **: p<0.01, ***: p<0.001.

### Remote representations engage hippocampal cell assemblies

The real-time system and the associated analyses presented above used all CA1 spikes above an amplitude threshold to assess the structure of hippocampal spatial representations. This approach provides more accurate decoding than using only spikes that can be confidently associated with single neurons^35,36,37^, but limits the conclusions that can be reached regarding single neuron activity. As memory retrieval is thought to engage the coordinated activity of ensembles of neurons^38^, we also performed spike sorting and analyzed the resulting putative single neuron data to determine whether and how ensembles were engaged during periods of remote representation.

Our results demonstrate engagement of coordinated ensembles of single neurons during remote spatial representations. We often found that multiple single neurons had place fields (increased firing rate) in the target region and that these individual neurons were active around the time of target representation detection (Fig. 6a). These included neurons that showed a pronounced increase in activity at detection of the reward-triggering representation (cells 1 and 2) and others that were active before, during, and after the detection of the reward-triggering representation (cells 3 and 4). We quantified this engagement by identifying coordinated activity of groups of neurons (cell assemblies^39^; see Methods) and segregating them into two groups based on whether or not their activity represented the target location during the exploration period of each session (target and non-target assemblies; Fig. 6b,c). We then computed the activity of these assemblies at the times when remote representations were expressed.

**Figure 6.**
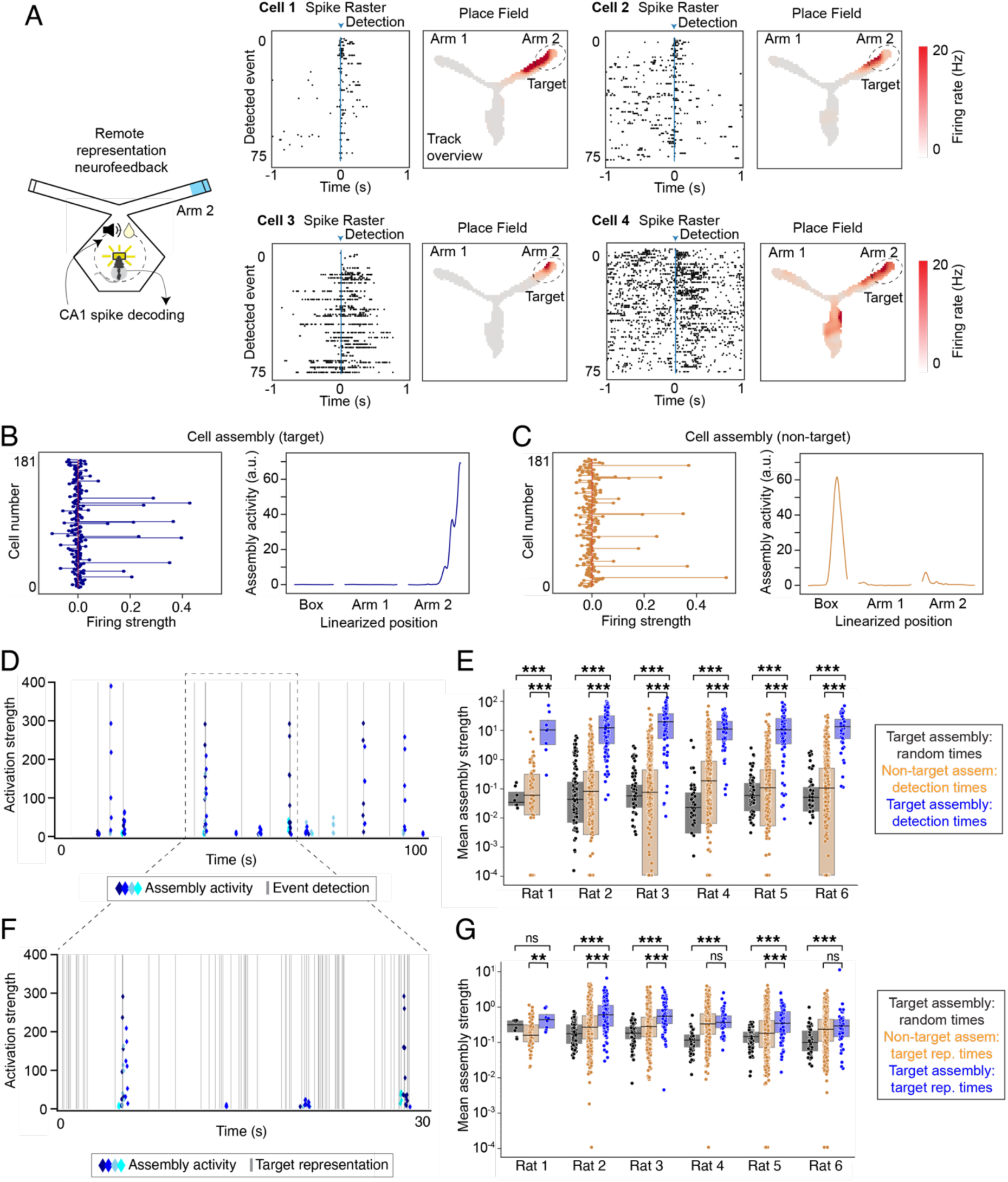
Cell assemblies are activated at the time of remote representations. **(A)** Individual neurons active at remote representation detection times. For each cell, left: raster plot showing spikes surrounding each detected remote representation (blue arrow); right: place field computed during exploration phase. **(B)** Target cell assembly. Left: individual cell weights; right: combined location-specific assembly activity. **(C)** Same plot as **(B)** for a nontarget cell assembly. **(D)** Example of activation strength for four target assemblies at the time of remote representation detection. Blue diamonds show activity of each target assembly and grey lines indicate detection times. **(E)** Target assembly activity at time of remote representation detection (blue) compared to random times (black) and to non-target assemblies at detection times (orange) across all sessions. **(F)** Zoom-in on plot from **(D)** showing target representation times (grey lines) and assembly activity (blue diamonds). **(G)** Target assembly activity at time of remote representation outside of detection events (blue) compared to random times (black) and to non-target assemblies at remote representation times (orange) across all sessions. ***: p < 0.001, Mann-Whitney test. In **(E)** and **(G)**, for plotting only, any values less than 1e-4 were set to 1e-4.

We first focused on times when the target representation was detected (and the reward cue was triggered) during the feedback period of the neurofeedback sessions. During these periods, when the animal was in the box, far from the actual target location, we nevertheless found strong and specific activation of target-representing assemblies: these assemblies were about 90 times more active at detection times compared to random times (Fig. 6d,e, each rat p<0.001, Mann-Whitney test) and were about 25 times more active than non-target assemblies (Fig. 6d,e, each rat p<0.001, Mann-Whitney test). In addition, animals frequently generated representations of the target location outside the times of rewarded events. During these periods, target assemblies were about 4 times more active than at random times (Fig. 6f,g, 5 of 6 rats p<0.001, 1 rat, p=0.16, Mann-Whitney test; grouped analysis of all rats, p = 3.2e-27, LME) and were about 1.6 times more active than non-target assemblies at these same times (Fig. 6g, 4 of 6 rats, p<0.001, 2 rats: p=0.4, 0.44; grouped analysis, p = 9.5e-18, LME). We note that target assembly activity was not detected during all times of remote representation; this is likely because our system used clusterless decoding (decoding without cell clustering using all spikes above a voltage threshold), which incorporates many spikes that are not clustered into single units during spike sorting^40^.

Finally, to verify that each detection event engaged multiple neurons (illustrated in an example, Fig. S6a), we counted the number of high-strength cells in target assemblies that were active before each target detection event (Fig. S6b, Methods). Across all sessions, rats had an average of 1.8 – 8 cells active immediately preceding each detection event, which corresponded to 10 – 15% of high assembly strength cells (Fig. S6c). This is comparable to the number of neurons expressing remote representations during movement periods as reported in previous papers^22,23^, and given that we are recording a tiny fraction of the total number of neurons in the hippocampus, suggests that many neurons are engaged during each of these remote representation events.

We then asked whether there were consistent changes in spatial representations associated with the generation of remote representation. Previous work has shown that the firing fields of place cells can move (remap) within the same environment when the location of reward changes^41^. Because rats in this task received reward at the center port while representing a remote location, and because the animals were activating representations of a remote location, it could have been the case that there were consistent reallocations of activity from the remote location to the center. This was not consistently true across animals, but there was some evidence for remapping in two of the six animals (Fig. S6d).

### Brain state during remote representation events

Previous work has identified remote spatial representations in the hippocampus in the context of two distinct physiological states. First, remote representations can be expressed during sharp-wave ripple (SWR) events. During waking, SWRs occur primarily during immobility, and spiking during SWRs is most strongly associated with memory consolidation processes^24,25,27,42^. Second, remote representations can be expressed in moving animals in the context of the ∼8 Hz theta rhythm, where late phases of theta are most strongly associated with remote activity^21–23,43^. Remote activity during theta is hypothesized to contribute to memory retrieval in the context of deliberative decision making^23,44^. We therefore asked whether the remote spatial representations occurred during SWRs, theta, or in a different brain state^45^ (Fig. 7a).

**Figure 7.**
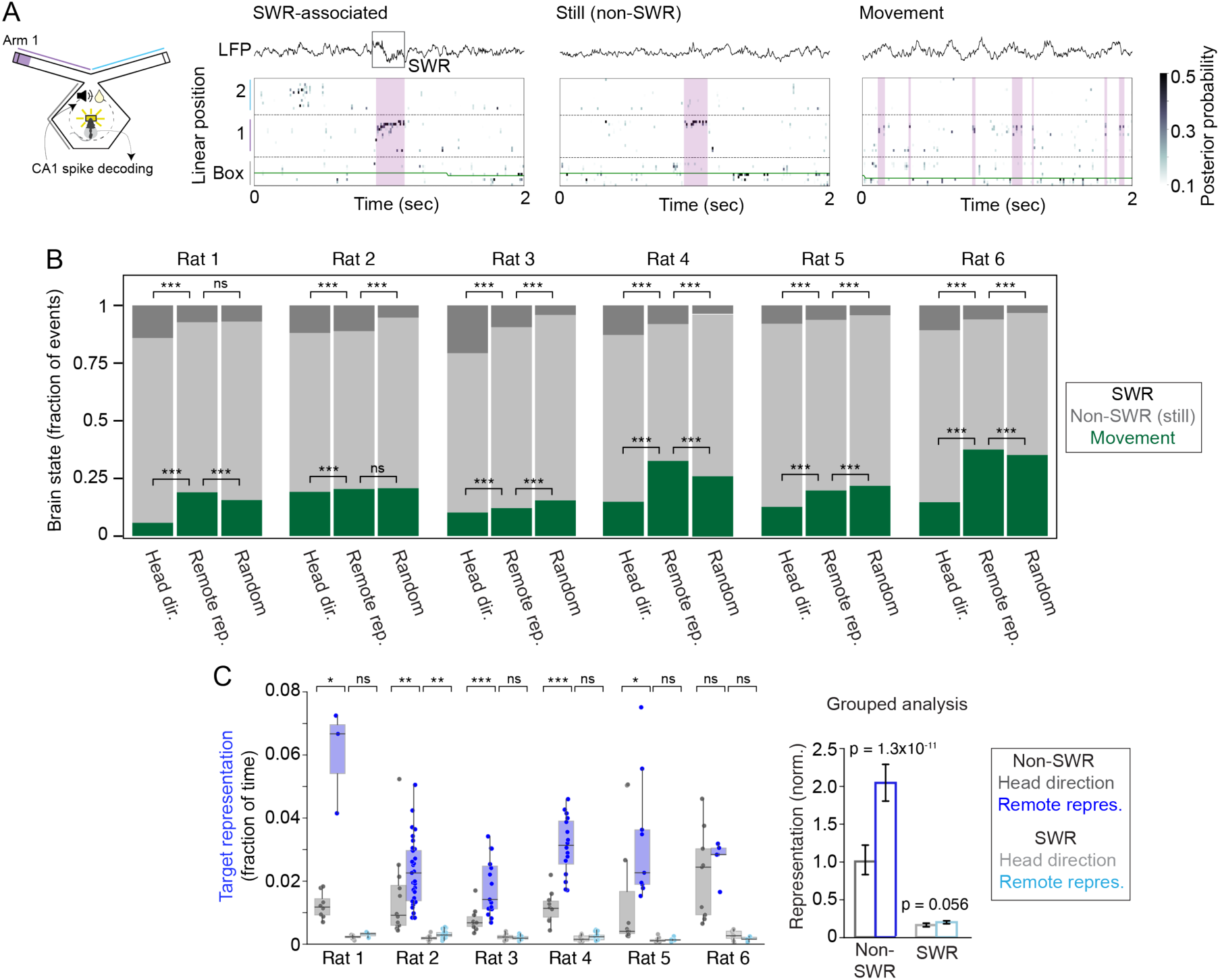
Brain state during remote representation. **(A)** Example remote representations during SWR, stillness outside of SWR, and movement. Left: schematic. Right: example plots; top: LFP trace, bottom: decoded linear position. **(B)** Summary of brain state during feedback periods. For each rat: remote representation during head direction feedback, remote representation during neurofeedback, and random times. ***: Chi-squared test and post-hoc z-test of proportions, p<0.001. **(C)** Change in target representation during SWR-associated and non-SWR times for neurofeedback sessions compared to head direction feedback sessions. Grouped analysis (LME) for 6 rats. *: p<0.05, Mann-Whitney test, **: p<0.01, ***: p<0.001.

Strikingly, remote representations were primarily expressed during stillness (head speed <4 cm/sec) but outside of clearly identifiable SWR events: only 10-20% of remote representation were associated with SWRs (Fig. 7b). Furthermore, target representations outside SWRs consistently increased in prevalence with neurofeedback (Fig. 7c, p < 0.05 in 5 of 6 individual rats, Mann-Whitney; grouped analysis: p = 1.3e-11, LME), while representations associated with SWRs did not consistently increase across animals (Fig. 7c, p < 0.05 in 1 of 6 rats, Mann-Whitney; grouped: p = 0.056, LME). Those remote representations that were expressed during movement showed characteristics similar to those seen in previous reports^3^. These representations were associated with specific theta phases in all rats (p < 0.0001 for each rat, Rayleigh test, Fig. S7a), and in 5 of 6 rats, remote representations were more prevalent in late theta phases.

We also compared the brain states during which remote representations were generated across the head direction and neurofeedback sessions. During neurofeedback, remote representations were more likely to occur during movement and less likely to occur during SWRs as compared to head direction feedback (Fig. 7b, Chi-square test and post-hoc Z-test of proportions for movement fraction and SWR fraction in each rat, p<0.001). This suggests that remote representations during neurofeedback occurred in more active and engaged states compared to those seen during head direction feedback. Within neurofeedback sessions, we compared brain states during remote representations to random times at the center reward port. Of note, SWRs were more frequent during times of remote representation compared to random times in 5 of 6 rats (Fig. 7b, Chi-square test and post-hoc Z-test of proportions for SWR fraction in each rat, p<0.001).

Given the surprising observation that most remote events occurred during stillness but outside of SWRs, we then asked whether there was a specific LFP signature associated with these events, as there is for immobility-related place activity^46^. We were not able to not identify such a signature, although some rats had increased power >100 Hz, potentially reflecting high gamma activity or spiking (Fig. S7b). Indeed, and as expected, multiunit spiking activity peaked at the time of remote representations (Fig. S7c). Finally, we also examined the relative prevalence, across brain states, of representations that jumped to the target location as compared to representations that included the intermediate locations along a trajectory from the animal to the target. We found both jump and trajectory representations across moving, still, and SWR periods and no consistent effects of prevalence across animals (Fig. S7d).

## DISCUSSION

We developed a closed-loop neurofeedback system for hippocampal representations and used this system to reward rats for generating specific remote spatial representations. Although no specific retrieval cues were presented, rats learned to activate specific representations corresponding to experimenter-chosen spatial locations. These representations typically jumped directly to the target region. Rats were also able to generate different representations at different times in response to changes in the target location, demonstrating flexibility. These remote representations engaged one or more cell assemblies and multiple neurons within each assembly, consistent with the reactivation of a population of neurons representing the remote location. Our work establishes a model for studying how the brain can deliberately reinstate representations related to previous experience in the absence of cues and specific behavioral outputs.

Studying memory retrieval as a distinct process requires distinguishing retrieval related activity from both the behaviors that retrieval can drive as well as the cues that can drive retrieval. Our work builds on previous brain-machine interface (BMI) approaches to make this possible. BMI systems can be used to drive animals to generate patterns of activity in the absence of associated behavior^47–49^, potentially enabling separation of retrieval from action. At the same time, previous approaches, including a recent demonstration of volitional control in the hippocampus^18^, use continuous sensory feedback to drive incremental changes in neural activity patterns that allow an animal to achieve a desired goal.

As our goal was to remove the potential confounds of pattern-specific sensory inputs, we instead required that animals remain near the center port of the maze (from which the experimenter-selected target region was not visible) and rewarded the animal for generating a representation of the target location. This approach had an additional critical benefit: it meant that animals could directly engage the representation of the target region without the requirement that they activate a series of intermediate representations. In the context of space, these would be the representations of the position between the animals’ actual location and the target presentation.

Our animals took advantage of that feature, generating representations that most often “jumped” to the target area without activating representations of intermediate locations. This enables study of a key aspect of memory: the ability to mentally teleport to places or events distant in space and time. The fact that the animals generated these distant events also argues strongly against a simple sensory explanation for their ability to activate the associated representations. From the neurofeedback location, the animal could see only the base of each of the two possible target arms. If visual cues were assisting the animal in generating a representation of the arm end (target), we would expect to see representation of the very beginning of the arm preceding representation of the target. However, we rarely observed these activity patterns immediately before target representations. Related to this, in the control experiment when the rat was rewarded for turning its head towards the target arm, the decoded representation remained in the box at the animal’s current location (Fig. S3d), further indicating that the remote representations generated during neurofeedback were not tied to the animal’s gaze. If cues in the visual field were driving remote representation, we would expect head turns towards the target to trigger remote representation as was recently reported in chickadees^50^.

Our work also provides insights into the brain states and activity patterns that can support the generation of remote representations consistent with memory retrieval. Most remote representations in this task occurred during stillness but did not overlap with SWRs, and remote representations that did occur during SWRs were not enriched despite many days of training. Thus, while it is possible to train animals to increase the rate of SWR generation^51^, these findings suggest that representations within SWRs are less amenable to control. This possibility is consistent with the idea that SWRs are critical for “offline” memory consolidation and updating functions rather than “online” retrieval to inform immediate decisions^25,24,27^.

We also found that some of the events occurred during movement, and consistent with previous observations^21,23,43^, these remote events were most often not in the trough of the theta rhythm where coding for the animal’s actual position is prevalent, but instead in the later phases of theta. This provides further support for the notion that specific theta phases are most conducive to the generation of non-local representations.

At the same time, most events were seen during periods of stillness and were not associated with an obvious local field potential signature. This state has also been associated with immobility-related place cell activity^46,52,53^, and our results indicate that in rats, the state is also consistent with a more “generative” function associated with remote representations^45^. Specifically, the lack of movement and associated sensory input may be conducive to the generation of internally driven patterns of activity associated with specific past experiences.

The specific mechanisms that support the generation of these remote representations are unknown, but previous studies of remote hippocampal representations in the context of SWRs^24,27^ and movement^21,28^ highlight the engagement of multiple cortical regions including the prefrontal cortex before, during, and after remote hippocampal events. It seems plausible that related networks are engaged when animals deliberately activate specific remote representations, but the mechanisms that might differentiate more automatic from more deliberate activation are not understood. More broadly, the generation of patterns of activity associated with past experience is central to the retrieval of memories^54^ and to related cognitive functions like planning and imagination^44^. The ability to train animals to generate such patterns in the absence of specific cues and specific behavioral outputs thereby provides a potentially powerful tool for understanding how the brain can support these remarkable functions.

## ACKNOWLEDGMENTS

We thank all members of Loren Frank’s lab for thoughtful feedback throughout the project. We thank Vanessa Bender, Massimo Scanziani, and Andrew M Lee for feedback on the manuscript. This work was supported by grants from the National Institutes of Health (F32MH123003 to MEC), the Simons Foundation (grant 542981 to LMF) and the Howard Hughes Medical Institute (grant to LMF).

## AUTHOR CONTRIBUTIONS

MEC designed and performed the experiments, analyzed the data, and wrote the manuscript. AKG helped design the experiments. JC, ELD, TTKN, DLF, KW, and XD contributed to the real-time decoding software. KW and BS helped perform the experiments. UTE and CK helped supervise the project. LMF supervised the project, helped design the experiments and analyses, and wrote the manuscript.

## DECLARATION OF INTERESTS

The authors declare no competing interests.

## SUPPLEMENTAL INFORMATION

**Figure S1.**
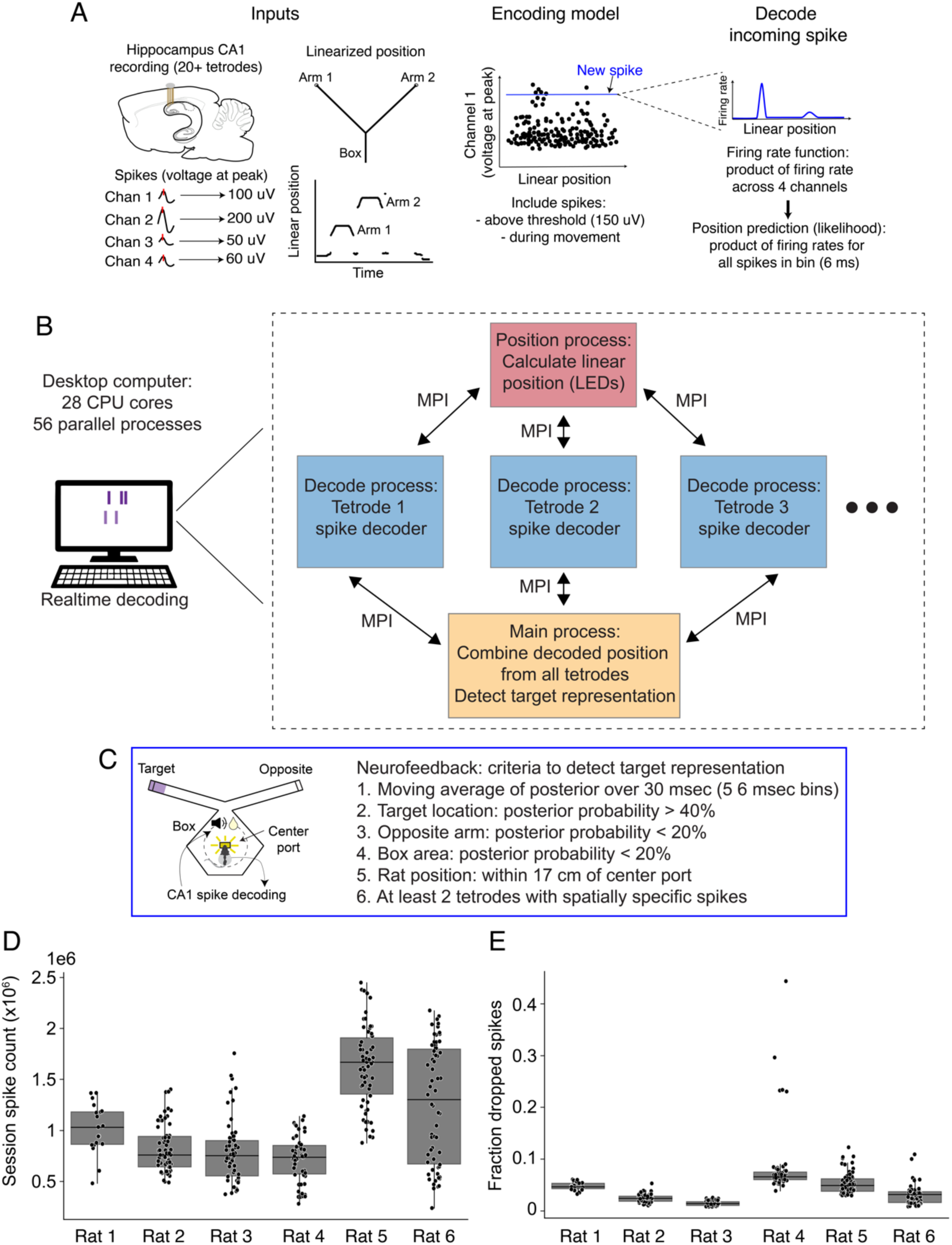
(**A**) Schematic of clusterless decoding algorithm. (**B**) Software architecture for real-time decoder implementation. MPI: message passing interface. (**C)** Criteria for detection of remote representation to trigger reward cue. (**D**) Total recording session spike counts for each rat. **(E**) Fraction dropped spikes for each rat (not decoded because of decoder latency).

**Figure S2.**
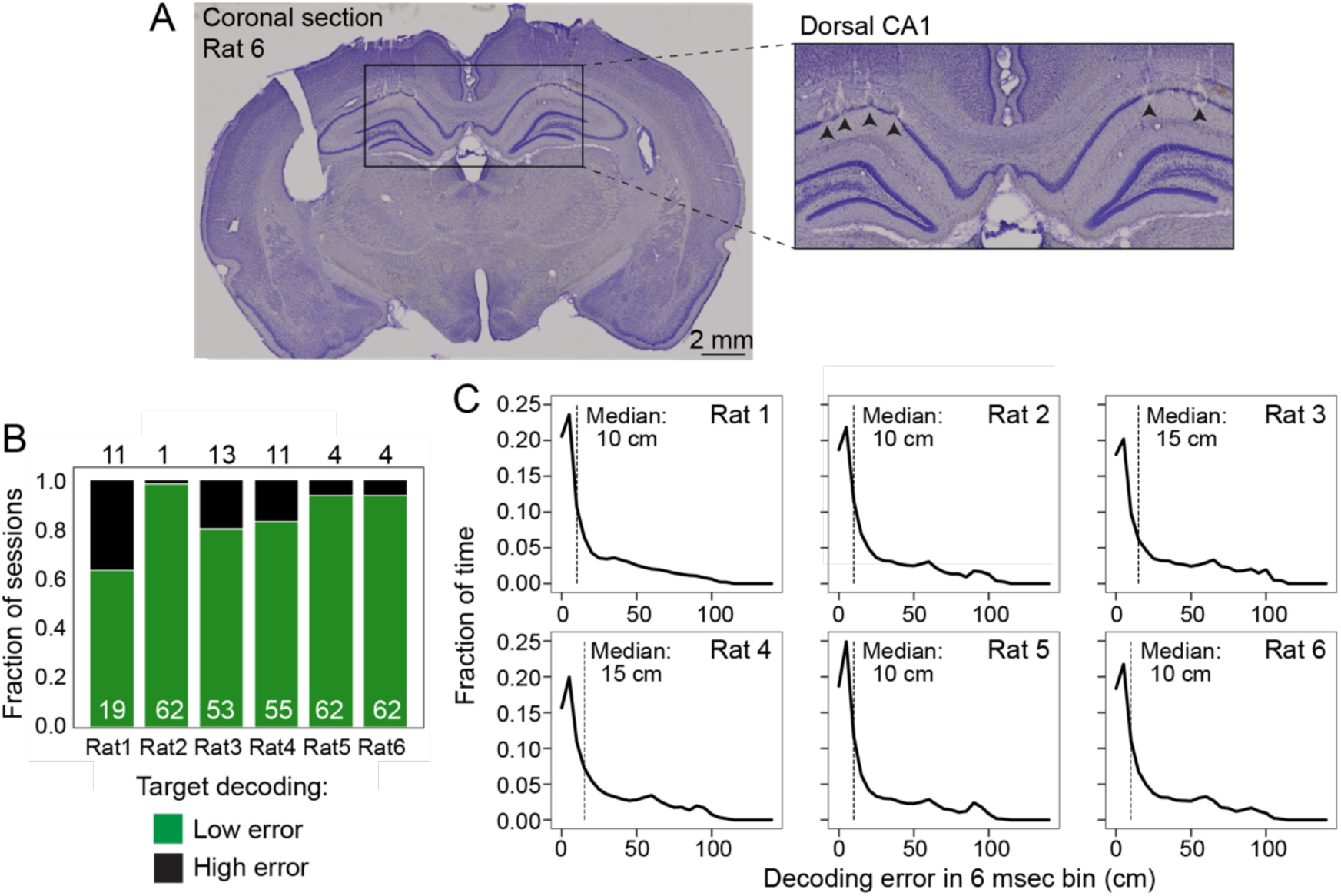
(**A**) Histology (Nissl stain) showing recording tetrodes located in dorsal CA1 of hippocampus. (**B**) Fraction of recording sessions for each rat with low decoding error vs. high decoding error (low: <35% decoding error in target region). (**C**) Decoding error during movement. Histogram of error (distance between rat’s real position and decoded mental position) for each 6 msec time bin while the rat was moving (>4cm/sec). Median is marked with dashed vertical line.

**Figure S3.**
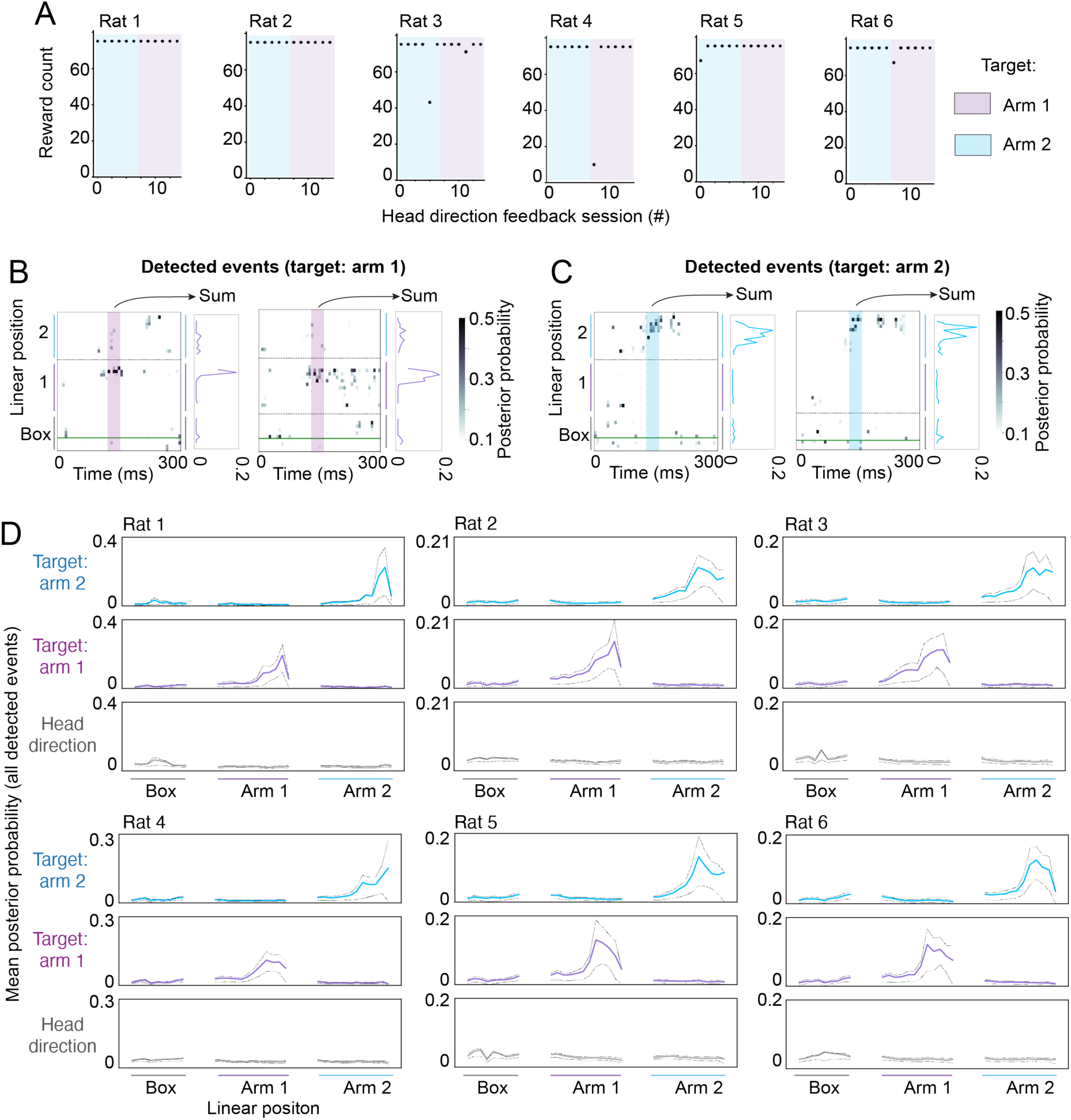
**(A)** Reward counts for all head direction feedback sessions. **(B)** The rats received reward when the decoded position matched the target location. Examples of rewarded representations for arm 1 (**B**) and arm 2 (**C**). Shaded areas contain time bins that contribute to detection (30 msec). The linearized sum of the representation is shown for each example (right plot). **(D)** Hippocampal representation averaged across all detected events. Organized by target region: arm 2 neurofeedback (blue), arm 1 neurofeedback (purple), or head direction feedback (grey). Posterior probability was averaged over 30 msec that triggered detection (as show in in (**B**)). Dashed lines represent 75% confidence interval.

**Figure S4.**
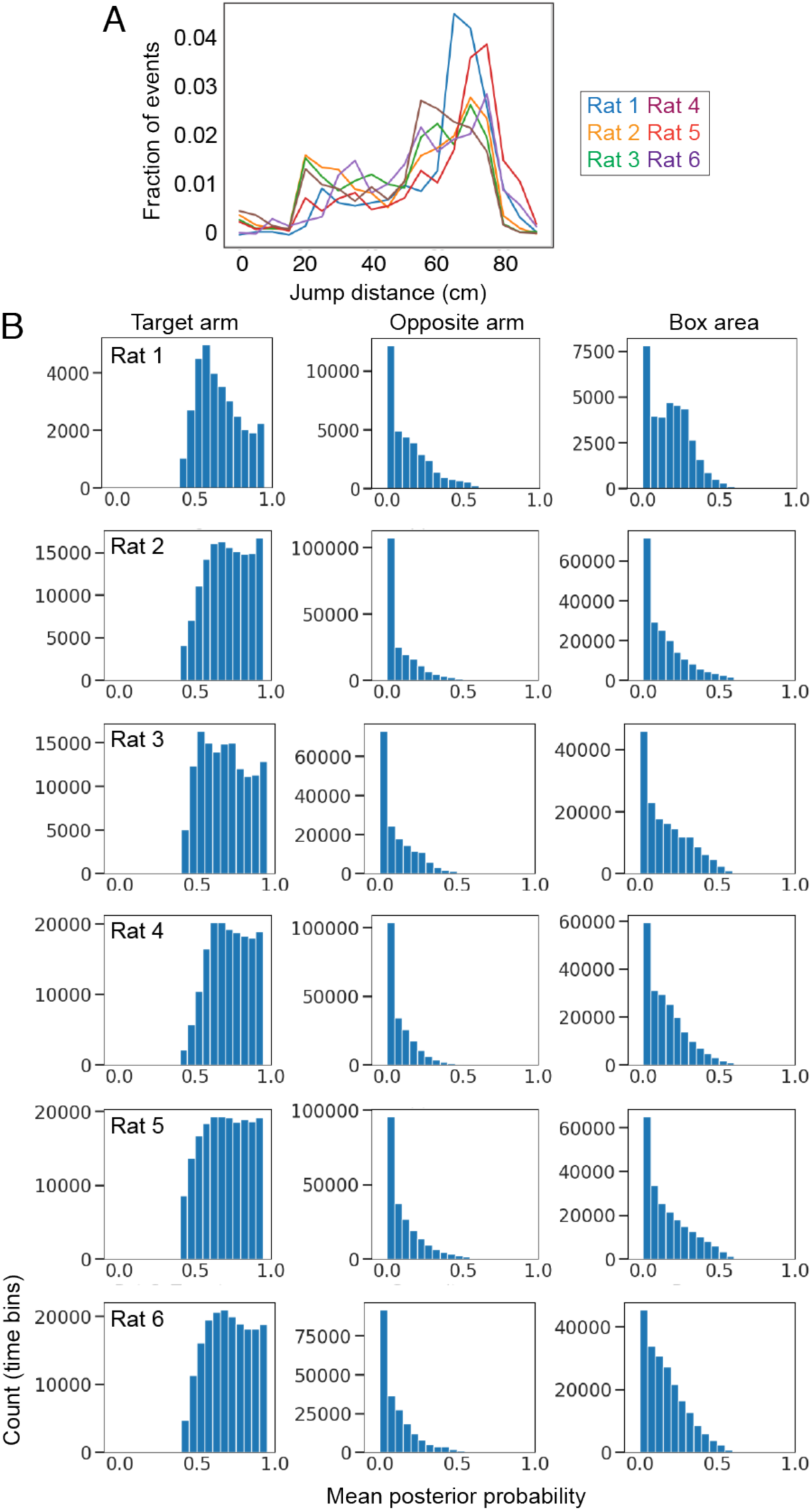
(**A**) Distance between rat’s actual position and nearest representation in target arm for detected events. Includes 90 msec before detection. (**B**) Decoded posterior probability at target representation times. This shows that for each rat, nearly all times of target representation had at least 50% representation in the target arm and much less representation of either the opposite arm or the local area (box).

**Figure S5.**
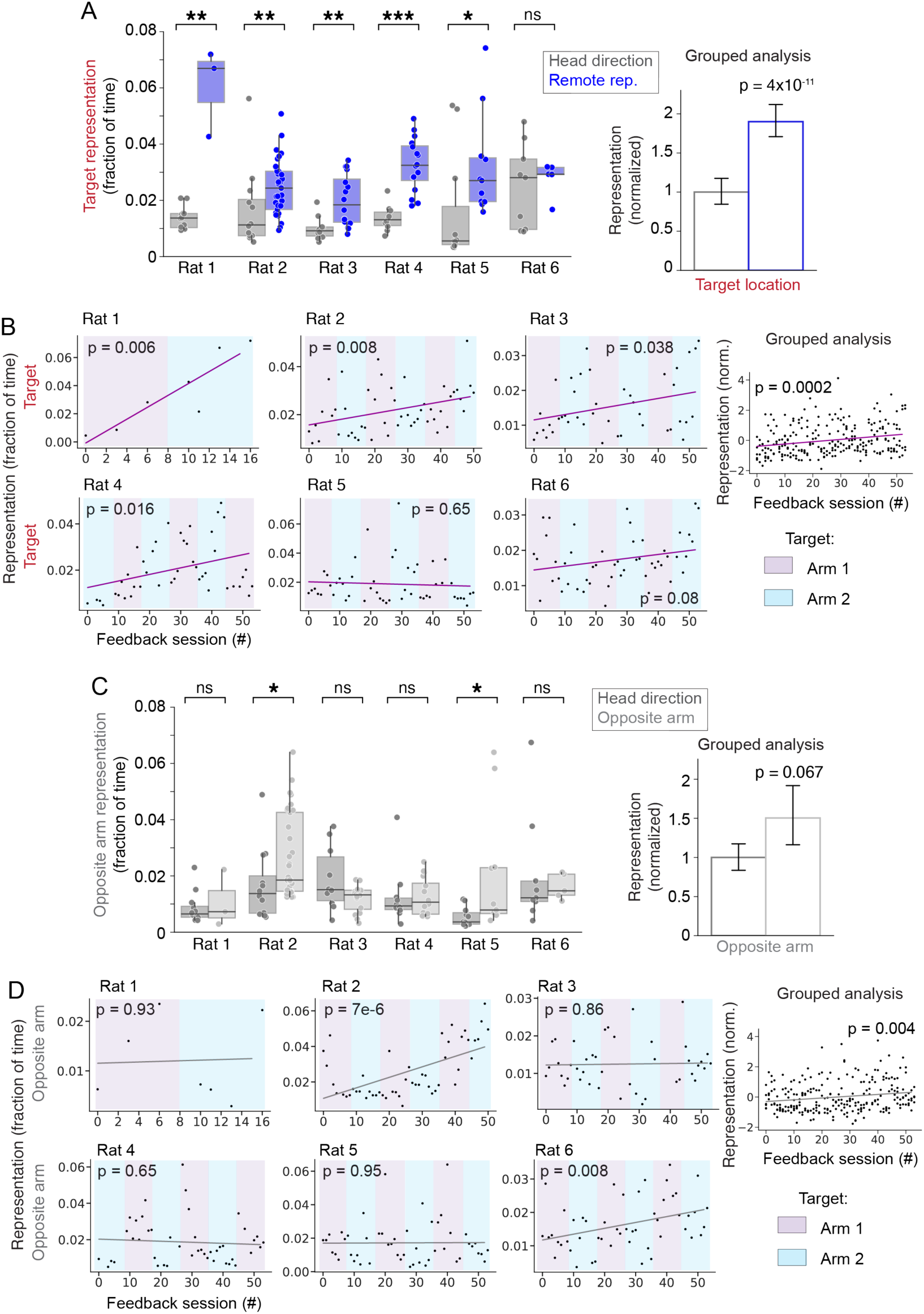
**(A)** Same plots as in Figure 2C with triggering representation removed. **(B)** Same plots as in Figure 2D with triggering representation removed. *: p<0.05, Mann-Whitney test, **: p<0.01, ***: p<0.001. **(C)** Same plots as in Figure 2C showing representation of opposite arm end (non-target). **(D)** Same plots as in Figure 2D showing representation of opposite arm end. *: p<0.05, Mann-Whitney test.

**Figure S6.**
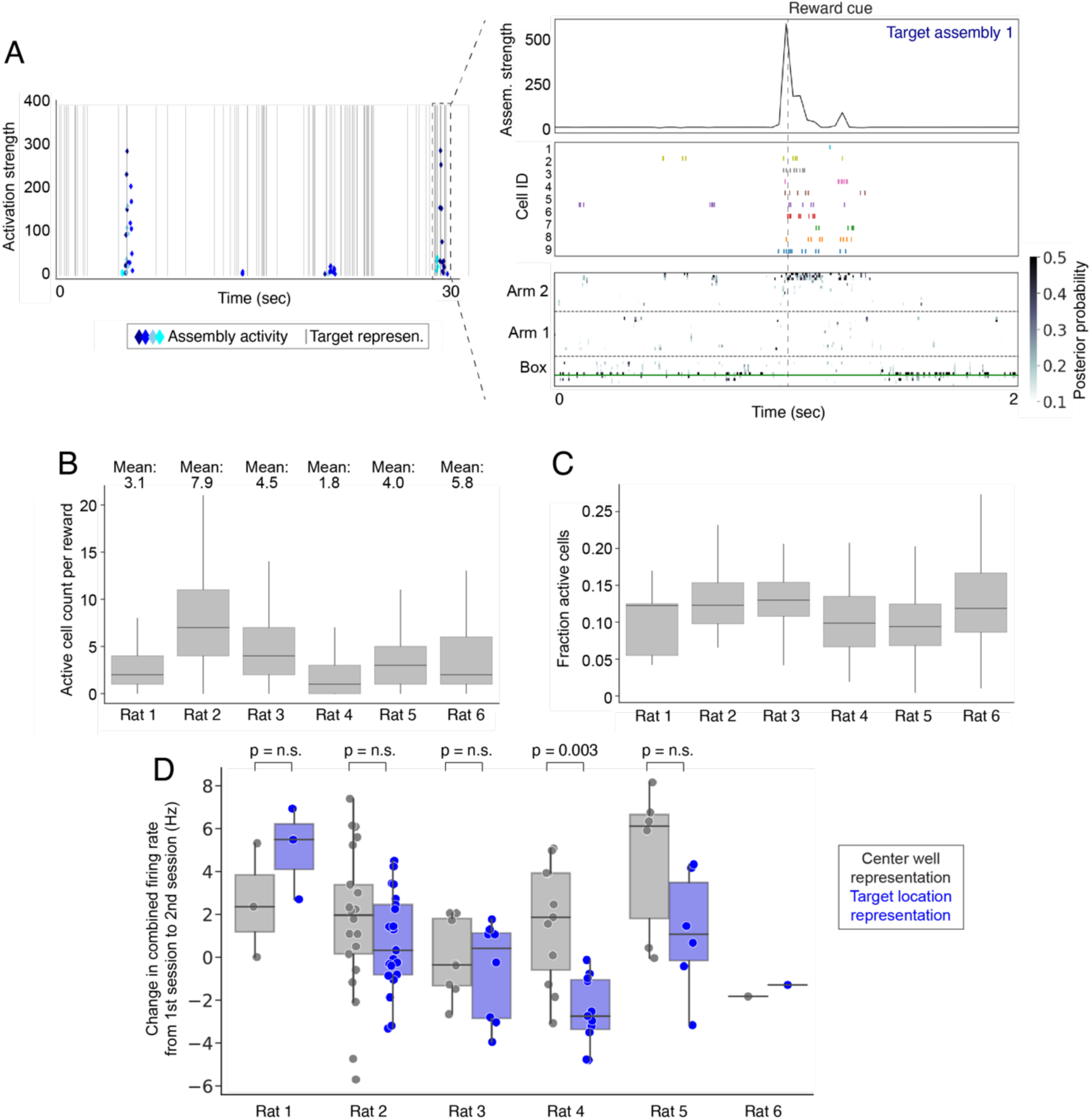
**(A)** Left: same panel from Figure 3F, showing target assembly activity (blue diamonds) and remote representation (grey lines). Right: zoom in on a detected target representation. During this representation, target assembly activity is high (top) and there are spikes from several high-strength cells within this assembly during the representation (middle). Clusterless decoding showing the target representation at the end of arm 2 (bottom). **(B)** We counted the number of high-strength cells with spiking activity immediately before target representations and found that, on average, all rats have more than one spiking cell (within 100 ms). **(C)** This plot shows the fraction of high-strength cells active before each target representation. The fraction is similar across rats showing that much of the variability in cell count in **(B)** is the result of variations in the number of high-strength cells across rats, an indicator of recording quality. **(D)** Population-level remapping analysis. Following high reward sessions, change in representation of neurofeedback location at center reward port (gray) and target region (blue) are shown for the subsequent session. There was no consistent pattern of change observed across animals (2 rats center port > target, 3 rats target > center port). Rat 4 had a significantly larger increase in representation for the center port compared to the target region (Mann-Whitney, p=0.003).

**Figure S7.**
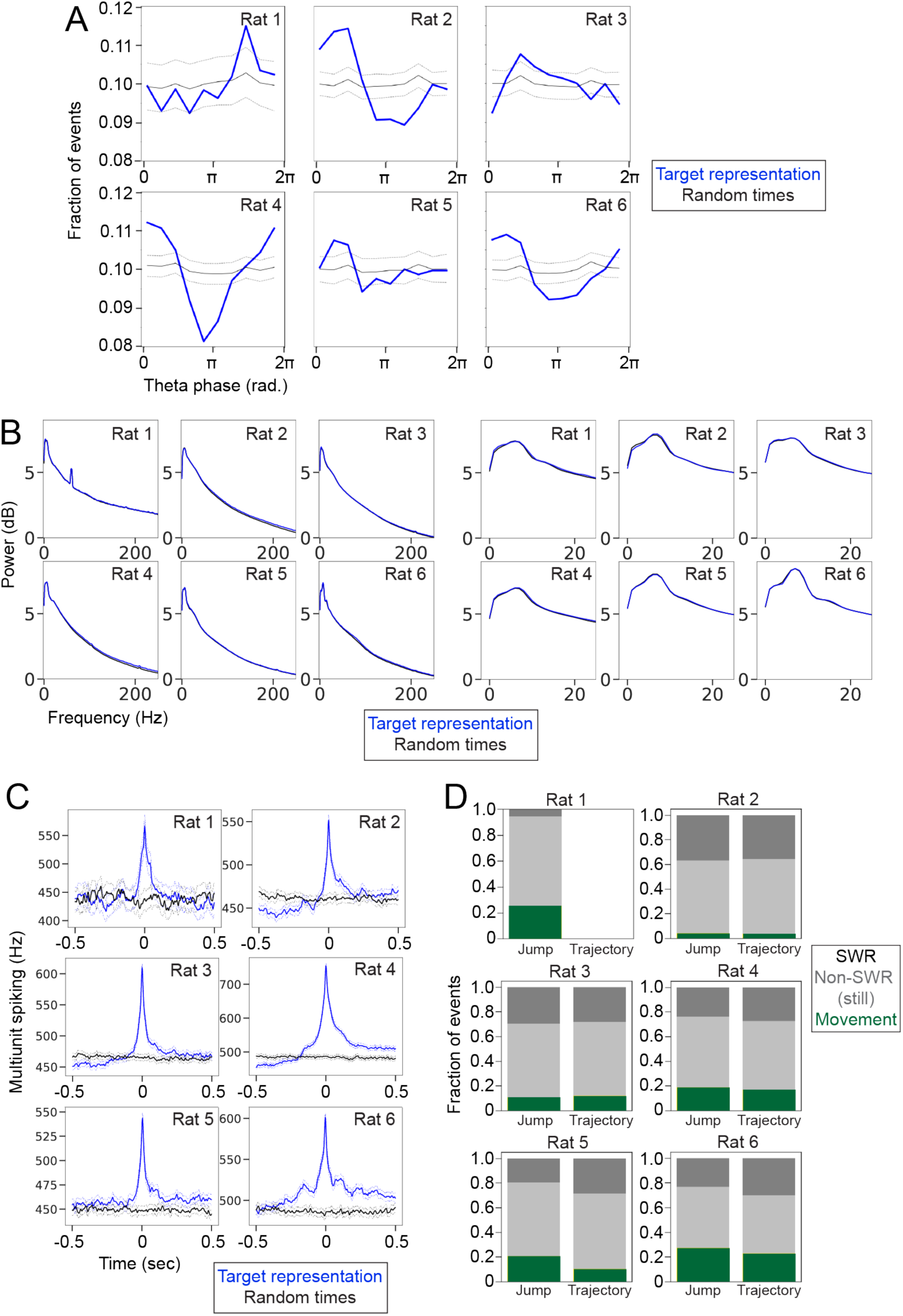
**(A)** Remote representation times (blue) are modulated by theta phase. Dashed line are 95% confidence interval for theta phase of random times. All possible theta phase values were grouped into 10 equal bins and the fraction of remote representation events in each bin is plotted. **(B)** LFP spectrogram of remote representation times. Left: tetrode in CA1 cell layer (referenced), right: tetrode above cell layer (unreferenced). Theta power is most accurately measured above the cell layer with an unreferenced tetrode, and so right plot is zoomed in to low frequency range. **(C)** Multiunit spiking at times of remote representation. Blue: remote representation, black: random times. Dashed lines: 95% confidence intervals of the mean. **(D)** Brain state associated with different types of remote representations (either jump or trajectory representations). There were no consistent differences across animals in the brain state associated with Jump or Trajectory representations.

## STAR METHODS

### RESOURCE AVAILABILITY

#### Materials Availability

Not applicable.

#### Data and Code Availability

Raw data will be available on DANDI.

Analysis code is available on a GitHub repository called: https://github.com/orgs/LorenFrankLab/hippocampus_content_neurofeedback

Realtime spatial decoder code is available on GitHub repositories called: Current version of code:

https://github.com/orgs/LorenFrankLab/realtime_decoder

Version of code used in the manuscript:

https://github.com/MichaelCoulter/realtime_decoder_v0

### EXPERIMENTAL MODEL AND SUBJECT DETAILS

Adult wild type Long Evans rats were obtained from Charles River Labs. Housed at UCSF following all IACUC guidelines. All surgical procedures done following IACUC guidelines.

### METHOD DETAILS

#### Behavioral training

Adult wild type Long Evans rats (6-9 months of age) were food restricted to 85% of their free body weight. Before neural implant surgery, rats were trained to nosepoke in reward ports to receive a milk reward on a linear track. Rats were then exposed to the task environment and trained in the task structure, in which they had to visit lit reward ports to receive a milk reward. The task environment consisted of a central area with a milk reward port and two attached arms with a reward port at each arm end. The task structure consisted of two phases during 40-45 minute recording sessions. In the first 10-15 minutes (exploration phase), the rat explored the Y-shaped environment (center box area, arm 1, and arm 2) and received milk reward by nosepoking in each lit reward port (300 uL of sweetened condensed milk was delivered automatically via the Trodes recording software). The light cues directed the rat to visit each arm 12 times and return to the center after each arm visit (arm 1 and arm 2 visits were randomly ordered). In the second phase (operant conditioning), lasting 20-30 minutes, the rat could only receive reward at the center port. During the conditioning phase, a sound cue was played and then a milk reward was available at the center port. In pretraining, the rat learned the association between the sound cue and reward: when the sound cue played, the rat had to nosepoke within 3 seconds to receive reward. During initial training the sound cue occurred randomly (centered on intervals of 5-30 seconds). For head direction feedback, the sound cue was triggered when the rat turned its head in a specific direction For remote representation neurofeedback, the sound cue was triggered by online detection of a specific remote hippocampal representation (as in pretraining, the rat had to nosepoke within 3 seconds of the sound cue to receive the milk reward).

#### Neural implant

Rats that completed the pretraining were surgically implanted with a Microdrive containing 64 individually moveable tetrodes targeting dorsal hippocampus^55^. This surgery was performed as described previously^25^ and in compliance with the UCSF IACUC animal safely protocols. Following surgery, tetrodes were lowered to the dorsal CA1 pyramidal cell layer over 2-3 weeks.

#### Clusterless decoding of hippocampal representation

Once tetrodes were lowered to the pyramidal cell layer, hippocampal spatial representations were decoded from recordings using a decoding algorithm described previously^35^. To decode position from action potential firing (spiking) of CA1 neurons, this algorithm builds a marked point-process model to relate all CA1 spikes (above a set threshold) to the rat’s position and then applies this model to estimate the position of each new spike. Position estimates for all spikes in a small time window (6 msec) are combined to generate a prediction of the spatial position represented in hippocampus. A key feature of this decoding algorithm is that it does not require spike sorting (“clusterless”).

#### Real-time spatial decoding

The clusterless decoding algorithm was implemented for fast operation using a desktop computer with 28 cores and parallelization of the decoding algorithm using message passing interface (MPI). During each recording session, the system decoded ∼1 million spikes. With a decoding latency of 30 msec, 90-99% of incoming spikes were decoded. The details of the real-time decoder are described in an accompanying manuscript^32^.

#### Head direction feedback

After surgery recovery and tetrode adjusting, each rat started with 4 days of head direction conditioning. Each training day included 3 recording sessions. The recording sessions followed the task structure described above. During the feedback part of the task, the sound cue and reward was triggered by the rat’s head direction. Reward cue was triggered by a head turn to either 30 degrees to left (arm 1) or to the right (arm 2) when facing forward towards the base of the outer arms. The angular accuracy required to trigger the reward cue increased over the first 25 rewards of each feedback session (from +/-20 degrees to +/-3 degrees). The feedback phase lasted for 30 minutes or until the rat received 75 rewards, whichever occurred first. For the first 2 days, the target direction was pointing towards one arm and then the target switched to the opposite arm for the next 2 days.

#### Remote representation neurofeedback

Following head direction feedback, the rat switched to remote representation neurofeedback. Each day had 3 recording sessions and the target location (either the distal 25 cm of arm 1 or arm 2) switched every 3 days. Based on recording quality rats were trained for 6 days (rat 1) or 18 days (rats 2-6). During the exploration phase at the beginning of each session, the recorded CA1 spikes were used to build an encoding model that associated spiking activity with spatial locations in the environment. Then, during the feedback phase, when the decoder detected a coherent representation of the target location, the sound cue was triggered and reward was available at the center port. The following requirements had to be met to trigger the reward cue: 30 msec running average of decoded position, >40% in target arm end, <20% in opposite arm, <20% in box area, at least 2 tetrodes with spatially specific spikes during 30 msec window, and rat physical distance to center port < 17cm. The feedback phase lasted for 30 minutes or until the rat received 75 rewards, whichever occurred first. In rat 1, only 1 tetrode with spatially specific spikes was required.

#### Data collection and processing

Hippocampal tetrode recordings were collected using SpikeGadgets hardware and Trodes recording software (sampling rate, 30 kHz). The rat’s physical position was recorded using a Manta camera (frame rate 30 Hz) and red and green LEDs attached to the headstage were tracked using Trodes. These LEDs allowed calculation of head direction. Raw electrical signal, position tracking, and reward port nosepokes were extracted from the Trodes recording file and converted to an NWB file (https://github.com/orgs/LorenFrankLab/rec_to_nwb). The NWB file was inserted into a database organized by the Spyglass package for reproducible analysis of neural data (https://github.com/orgs/LorenFrankLab/spyglass)^56^. All subsequent analysis was done on data within the database.

#### Real-time decoder output

The decoded mental position as estimated by the real-time decoder was saved each session in a separate recording file. This file included the rat’s actual position and decoded position for every 6 msec time bin and each detected reward event (head direction or remote representation).

#### Data inclusion

Recording sessions were included with high quality spatial decoding. High quality sessions were defined as: during the exploration period, when the rat was running outward from the box and in the end of the target arm, the posterior maximum of the decoded position was in the target arm in at least 65% of time bins.

#### Spike Sorting

The raw tetrode recording was band-pass filtered from 600-6000 Hz and spikes were detected and sorted using MountainSort4^40^. During subsequent curation, noise clusters and multiunit clusters were removed automatically. Manual inspection was used to merge clusters that were inappropriately split. The remaining clusters are considered “isolated single units.”

#### LFP analysis

For LFP analysis, the raw tetrode recordings were band-pass filtered from 0 – 400 Hz. Previously described methods were used to detect sharp wave ripples^25^. Theta phase was calculated from one tetrode in each rat located above the CA1 pyramidal cell layer (stratum oriens).

#### Cell assembly analysis

Cell assemblies were identified using ICA with methods described previously^39^. The spike times of all isolated single units for a single recording session were binned into 30 msec bins and then co-firing cell assemblies were detected.

#### Histology

Rat brains were fixed with 4% PFA, serially sectioned, and then treated with Nissl staining to visualize hippocampal cellular structure and electrolytic lesions (made after the experiment) at the tetrode recording sites.

## QUANTIFICATION AND STATISTICAL INFORMATION

A few analyses could not be performed on all recording sessions because of errors with data saving and preprocessing. The missing sessions are listed here.

Figures 1-5: no missing data

Figure 6: rat 1 (2 missing sessions), rat 2 (2 missing), rat 3 (1 missing), rat 4 (1 missing), rat 5 (4 missing), rat 6 (3 missing).

Figure 7: rat 1 (4 missing head direction, 2 missing neurofeedback), rat 2 (0 missing), rat 3 (1 missing neurofeedback), rat 4 (0 missing), rat 5 (4 missing neurofeedback), rat 6 (2 missing neurofeedback).

Figure 3b

Fraction of detected events in each of 4 representation categories: jump (remote representation confined to target location with or without center port), jump and arm base (subset of jump events with representation at the base of the target arm), medium trajectory (representation with significant linear regression covering at least 35 cm in target arm), long trajectory (representation with significant linear regression covering at least 45 cm in target arm). Definitions: Target location: 25 cm at the end of the arm. Base of arm: 20 cm at the base of the arm, closest to the box. For a representation to be counted at the arm base or target location, the maximum of the posterior was required to be within that location and the posterior mass in the location had to be >0.4.

Figure 3c

Fraction of detected events with remote representation within the rat’s visual field (first 5cm of target arm), versus representation without visual field representation.

Figure 3d

Fraction of detected events with strong representation in specific reward port location within target arm, versus fraction of events without representation at reward port.

Figure 4

Spatial representation was calculated by counting the number of 6 msec time bins while the rat was near the center reward port (within 17cm) and > 40% of the decoded mental position (posterior mass) matched the specified location. For target representation (left), the location was the farthest 25 cm of the target arm from the box area (“arm end”), for non-rewarded representation (right), the location was the closest 25 cm of target arms to the box area (“arm base”). To calculate the fraction of time, the number of time bins meeting these criteria was divided by the total number of time bins at the reward port during the feedback part of the session. To match the amount of reward for head direction and remote representation conditioning, only sessions when the rats received >90% of the maximum rewards were included. Each dot represented one session, and significance was calculated with the Mann-Whitney test. Grouped analysis: a linear mixed model was created using the values just described grouped by rat. The tested variable was session type: either head direction or remote representation.

*n*: Rat 1: Head direction: n=12 sessions, neurofeedback: n=3. Rat 2: Head direction: n=12, neurofeedback: n=33. Rat 3: Head direction: n=10, neurofeedback: n=16. Rat 4: Head direction: n=10, neurofeedback: n=16. Rat 5: Head direction: n=11, neurofeedback: n=11. Rat 6: Head direction: n=9, neurofeedback: n=5.

Figure 5

In each rat, linear regression was performed on the fraction of target representation for each neurofeedback session. Top panels show change in target representation over time and bottom panels show change in non-target representation. Grouped analysis: fraction of target representation was z-scored within each rat and then data from all rats was combined and linear regression was performed on the combined data.

*n*: Rat 1: n=7 sessions, Rat 2: n=50. Rat 3: n=42. Rat 4: n=44. Rat 5: n=50. Rat 6: n=51.

Figure 6a

Place field for each cell was calculated as the occupancy normalized firing rate in the 2D task environment. Spatial bins are 3cm x 3cm.

Figure 6b and 6c

Left plot shows the weights of each cell for an example cell assembly. Right plot shows occupancy-normalized assembly activity at each location in the task environment.

Figure 6e

For each session, the firing of assemblies that represented the target location (highest assembly strength for either box, arm 1 end, or arm 2 end, with strength > 100) was compared at times of representation detection (reward cue) to randomly selected times while the rat was at the center reward port. Target assembly at reward cue times was also compared to non-target assembly (all other assemblies) at reward cue times. Significance calculated with Mann-Whitney test.

*n*: Rat 1: target: n=8 cell assemblies, random: n=8, non-target: n=72. Rat 2: target: n=126, random: n=126, non-target: n=694. Rat 3: target: n=81, random: n=81, non-target: n=527. Rat 4: target: n=57, random: n=57, non-target: n=568. Rat 5: target: n=85, random: n=85, non-target: n=578. Rat 6: target: n=53, random: n=53, non-target: n=769.

Figure 6g

For each session, the firing of assemblies that represented the target location was compared at times of target representation (>40% posterior mass in target location) to randomly selected times while the rat was at the center reward port. Target assembly at target representation times was also compared to non-target assembly (all other assemblies) at target representation times. Significance calculated with Mann-Whitney test.

*n*: Rat 1: target: n=8 cell assemblies, random: n=8, non-target: n=72. Rat 2: target: n=126, random: n=126, non-target: n=694. Rat 3: target: n=81, random: n=81, non-target: n=527. Rat 4: target: n=57, random: n=57, non-target: n=568. Rat 5: target: n=85, random: n=85, non-target: n=578. Rat 6: target: n=53, random: n=53, non-target: n=769.

Figure 7b

Each recording session was divided into three brain states: movement (rat speed > 4cm/sec), stillness (rat speed < 4 cm/sec outside of SWRs) and SWR (still within detected SWR). Decoding time bins (6 msec) with target representation (>40% posterior mass in target location) were assigned to each state across all recording sessions for each rat. The bar plot shows the fraction of these assignments for each rat. For random times, 5000 time bins were randomly sampled from the time the rat was at the center reward port and then assigned to each of the three brain states.

Figure 7c

This is the same analysis as Figure 2c, except that times during SWRs have been separated from the remainder of the recording session. Fraction of target representation was calculated as described during SWRs times and non-SWRs times. Grouped analysis same as Figure 2c.

*n*: Rat 1: Head direction: n=8 sessions, neurofeedback: n=3. Rat 2: Head direction: n=12, neurofeedback: n=33. Rat 3: Head direction: n=10, neurofeedback: n=15. Rat 4: Head direction: n=10, neurofeedback: n=16. Rat 5: Head direction: n=11, neurofeedback: n=9. Rat 6: Head direction: n=9, neurofeedback: n=5.

Figure S1d

Total number of spikes detected by the real-time decoder was counted for each included recording session.

*n*: Rat 1: n=19 sessions, Rat 2: n=62. Rat 3: n=53. Rat 4: n=55. Rat 5: n=62. Rat 6: n=63.

Figure S1e

Dropped spikes were defined as spikes that were decoded with a latency >30 msec. Fraction dropped spikes: dropped spike count / total spike count for each session.

*n*: Rat 1: n=19 sessions, Rat 2: n=62. Rat 3: n=53. Rat 4: n=55. Rat 5: n=62. Rat 6: n=63.

Figure S2b

High quality sessions (“good”) were defined as: during the exploration period, when the rat was running outward from the box and in the end of the target arm, the posterior maximum of the decoded position was in the target arm in at least 65% of time bins.

Figure S2c

Decoding error was calculated as the difference between the rat’s actual linear position and the posterior maximum for each 6 msec time bin during movement. The decoding error for all included sessions was combined for each rat and the plot is a histogram of error distance. Dashed line is median for all movement bins.

*n*: Rat 1: n=19 sessions, Rat 2: n=62. Rat 3: n=53. Rat 4: n=55. Rat 5: n=62. Rat 6: n=63.

Figure S3d

The decoded mental position that triggered reward (30 msec) was averaged across all sessions of each type for each rat (head direction, arm 1 representation, or arm 2 representation). These 3 average representations are plotted for each rat. Dashed lines are 75% confidence intervals of the data.

Figure S4a

Jump distance within detected remote representations. Distance from the center reward port to the closest remote location within the target arm with high confidence remote representation (posterior mass > 0.4). This analysis was performed on the decoded position from 90 msec before to the time of detection for each remote representation.

Figure S4b

Posterior for all position bins was averaged across all time bins with >40% representation of the target location. The full posterior was split into 3 non-overlapping categories: target arm, opposite arm, and box area. Plots show histogram of the posterior probability for each category across all included time bins.

Figure S5a

This is the same plot as Figure 2c except that the remote representations that triggered reward are excluded.

*n*: Rat 1: Head direction: n=12 sessions, neurofeedback: n=3. Rat 2: Head direction: n=12, neurofeedback: n=33. Rat 3: Head direction: n=10, neurofeedback: n=16. Rat 4: Head direction: n=10, neurofeedback: n=16. Rat 5: Head direction: n=11, neurofeedback: n=11. Rat 6: Head direction: n=9, neurofeedback: n=5.

Figure S5b

This is the same plot as Figure 2d except that the remote representations that triggered reward are excluded.

*n*: Rat 1: n=7 sessions, Rat 2: n=50. Rat 3: n=42. Rat 4: n=44. Rat 5: n=50. Rat 6: n=51.

Figure S5c

Same plot as Figures 2c except for opposite (non-target) arm end.

*n*: Rat 1: Head direction: n=12 sessions, neurofeedback: n=3. Rat 2: Head direction: n=12, neurofeedback: n=33. Rat 3: Head direction: n=10, neurofeedback: n=16. Rat 4: Head direction: n=10, neurofeedback: n=16. Rat 5: Head direction: n=11, neurofeedback: n=11. Rat 6: Head direction: n=9, neurofeedback: n=5.

Figure S5d

Same plot as Figures 2d except for opposite (non-target) arm end.

*n*: Rat 1: n=7 sessions, Rat 2: n=50. Rat 3: n=42. Rat 4: n=44. Rat 5: n=50. Rat 6: n=51.

Figure S6b

Cell number active at reward times was calculated by taking the strongest cells from each target assembly (strength > 5 s.d. above mean) and counting how many of these cells had spikes in the 100 msec preceding each reward.

*n*: Rat 1: n=322 reward times, Rat 2: n=2999. Rat 3: n=1970. Rat 4: n=1737. Rat 5: n=2037. Rat 6: n=1176.

Figure S6c

For this plot, the number of active cells at reward times was divided by the total number of high strength cells from reward assemblies, thus giving a fraction of measured cells active before each reward.

*n*: Rat 1: n=322 reward times, Rat 2: n=2999. Rat 3: n=1970. Rat 4: n=1737. Rat 5: n=2037. Rat 6: n=1176.

Figure S6d

Spatial representation changes after high-reward neurofeedback sessions. All place cells active during exploration (task phase 1) were identified, and their place fields were summed to generate a population-level representation during exploration. The strength of the spatial representations (firing rate in Hz) at the center reward port and the remote target location were compared between high-reward sessions (>67 rewards) and the subsequent session.

Figure S7a

Theta phase during time bins of remote representation (>40% posterior mass in target location). Only movement times were considered (rat speed > 4cm/sec). To create null distribution (grey line with 95% confidence interval dashed lines) theta phase was calculated at random time bins matching the number of remote representation time bins. The random sampling was done 1000 times and the distribution was used to calculate the 95% confidence interval.

*n*: Rat 1: n=13822 representation times. Rat 2: n=42027. Rat 3: n=21747. Rat 4: n=76860. Rat 5: n=37556. Rat 6: n=71226.

Figure S7b

Power spectrum of time (1 sec) surrounding remote representation. Spectra were calculated for each remote representation time bin outside of SWRs when the rat was still (speed < 4cm/sec) at the center well and then averaged. As a control, power spectra were calculated for random times when the rat was at the center reward port. Left plots: average power spectrum calculated from tetrode within pyramidal cell layer. Right plots: average power spectrum from tetrode above cell layer (used for theta band analysis).

*n*: Rat 1: n=1071 representation times (1 sec non-overlapping bins, rat still, non-SWR). Rat 2: n=16032. Rat 3: n=13605. Rat 4: n=17164. Rat 5: n=15025. Rat 6: n=14241.

Figure S7c

Combined population spiking at time of remote representations (surrounding 1 sec). Combined spiking was calculated by summing spiking of all isolated single units in 10 msec time bins. As a control, random times were selected when the rat was at the central reward port. Dashed lines are 95% confidence intervals of the mean.

*n*: Rat 1: n=1840 representation times (1 sec non-overlapping bins). Rat 2: n=18211. Rat 3: n=22150. Rat 4: n=21021. Rat 5: n=23581. Rat 6: n=23092.

Figure S7d

Brain state distribution during different types of remote representations. Distribution between brain states (movement, stillness, and SWR, see Fig 7) for jump representations (remote representation confined to target location with or without center port) and trajectory representations (posterior with significant linear regression covering at least 35 cm in target arm).

## Notes

### Competing Interest Statement

The authors have declared no competing interest.

### Summary of Updates

This version of the manuscript includes new analysis showing that remote representations primarily jumped directly to the target location without activating representations of intermediate locations (Figure 3). This enables study of a key aspect of memory: the ability to mentally teleport to places or events distant in space and time.

